# Feasibility of Measuring Magnetic Resonance Elastography-derived Stiffness in Human Thoracic Aorta and Aortic Dissection Phantoms

**DOI:** 10.1101/2024.09.05.611548

**Authors:** Adnan Hirad, Faisal S. Fakhouri, Brian Raterman, Ronald Lakony, Maxwell Wang, Dakota Gonring, Baqir Kedwai, Arunark Kolipaka, Doran Mix

## Abstract

Type-B aor tic dissection (TBAD) represents a serious medical emergency with up to a 50% associated 5-year mortality caused by thoracic aorta, dissection-associated aneurysmal (DAA) degeneration, and rupture. Unfortunately, conventional size related diagnostic methods cannot distinguish high-risk DAAs that benefit from surgical intervention from stable DAAs. Our goal is to use DAA stiffness measured with magnetic resonance elastography (MRE) as a biomarker to distinguish high-risk DAAs from stable DAAs. This is a feasibility study using MRE to 1) fabricate human-like geometries TBAD phantoms with different stiffnesses. 2) measure stiffness in TBAD phantoms with rheometry and 3) demonstrate the first successful application of MRE to the thoracic aorta of a human volunteer. Aortic dissection phantoms with heterogenous wall stiffness demonstrated the correlation between MRE-derived stiffness and rheometric measured stiffness. A pilot scan was performed in a healthy volunteer to test the technique’s feasibility in the thoracic aorta.

## Introduction

Type-B Aortic Dissection (TBAD) represents a serious medical emergency with up to a 50% associated 5-year mortality (*1, 2*). Dissection-associated aneurysmal (DAA) degeneration and rupture cause long-term TBAD mortality, and some clinicians have advocated for early surgical intervention, but adoption of this practice has been controversial. While some high-risk criteria imaging criteria have been developed to predict future DAA, none of these criteria have been validated. Fundamentally, the aneurysmal dilation of the aorta derives from a mismatch between vessel biomechanical wall properties and pulsatile stresses (*3*). This question, in part, can be resolved by understanding the natural history of material properties of the dissected aortic wall and its interaction with dissection-specific hemodynamics.

Although the biomechanical properties of abdominal aortic aneurysms (AAA) have been extensively profiled, little evidence exists on the evolution of the material properties in DAA over time. AAAs usually evolve over decades while DAAs may occur within a more acute period of months to years following an acute dissection. Despite this time-scale difference, we can use the extensive biomechanical evidence from AAA to guide the investigation of factors contributing to DAA. In AAA, stiffening of the aortic wall leads to pathological wall stress distribution and predisposes towards aneurysmal degeneration and rupture(*4*). Mechanically, increased stiffness due to loss of elastic lamina and increased collagen cross-linking is a significant predictor of AAA growth. In aortic dissection (AD), the medial layer, which comprises the elastic component of the aorta, is disrupted, making the false lumen consequently stiffer.

To date, one of the greatest constraints in the study of human AD pathology is the limitation of *in vivo* models and their ability to reproduce the biomechanical properties of human disease. Researchers have reported various chemically-induced ADs in small animal models. Unfortunately, these models have markedly different biomechanical profiles compared to human tissue (*5*) and therefore, new tools that directly study the human aorta are needed.

Magnetic Resonance Elastography (MRE) has emerged as an essential tool for imaging tissue stiffness in the liver, heart, brain, and vascular tissue(*6*). In vascular tissue, MRE has been applied in AAA(*7*), where aneurysm growth has been correlated with aortic wall stiffness but not aneurysmal size, the current clinical standard(*8*). In MRE, tissue stiffness is based on direct measurement of micro displacements that are created by externally produced mechanical shear waves(*9*). Given the similarities in the biomechanical behavior of AAA and DAA, MRE could similarly be applied to AD pathology.

While MRE has been applied to study the material properties of the abdominal aorta, and AAA, its feasibility in the thoracic aorta and DAA remains unknown. To this end, this study aims to investigate 1) stiffness contrast in hydrogel phantoms with human-like dissection geometries by using MRE and 2) demonstrate the first successful application of MRE to the thoracic aorta of a human volunteer.

## Methods

### TBAD Phantom Fabrication

Four, two-lumen TBAD phantoms were created, each comprised of heterogeneous material properties consisting of two differing PVA-c concentrations that mimic increasing stiffness throughout a previously reported range of human aortic tissue properties (*10, 11*). Athree-part polylactic acid phantom mold assembly, Figure 1, was printed using a Raise3D Pro2 Series desktop printer (Raise3D, Lake Forest, CA, USA). Phantom molds were designed to mimic an aortic dissection. Each mold has two distinct lumens with total diameter of 30 mm and a total length of 200 mm. The phantom has a 5 mm vessel wall and a 3 mm thick dissection flap, consistent with human AD geometriesIn all models, the right sidewall was composed of 10% PVA to replicate the “true lumen” and the left sidewall would represent the false lumen with increased stiffness composed of 10% [control], 15%, 20% or 25% PVA. Three mold subassemblies A, B, C, as seen in Figure 1, were required for phantom creation. Assembly A was injected with a designated concentration of 10%, 15%, 20% or 25% by weight PVA-c to create the false lumen sidewall. Assembly A then underwent a 12-hour freeze at -20°C to solidify the PVA-c. In the next step, the frozen assembly A and an empty assembly B were assembled inside assembly C. 10% PVA-c was injected into empty assembly B to create both the true lumen sidewall and the dissection flap. Assembly C prevented leakage of PVA-c and provided apposition of assemblies A and B for optimal bond cross polymerization. The assembly underwent four freeze-thaw cycles from -20°C to 20°C for PVA-c cross polymerization. Phantoms were then stored in chlorinated 20°C water until MRE testing. All models were tested within one month of creation to prevent changes in the hydrogel material properties.

**Figure 1:**
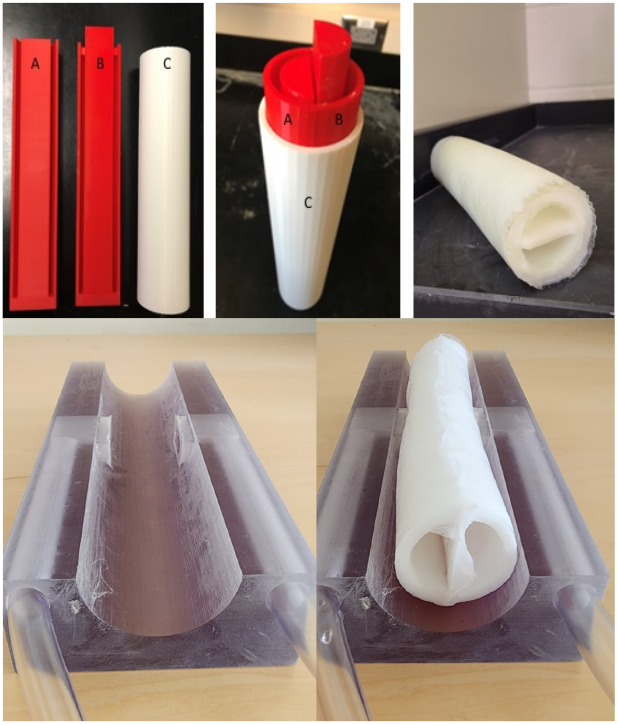
(***top left***) Shells A, B, and C, disassembled (***top middle***) Shells A, B, and C, assembled into the mold **(top *right*)** *Cyrogel* phantom removed from mold (***bottom left***) in house designed and built passive driver, with dual diaphragms to deliver vibrations to each side of the simultaneously. phantom separately but (***bottom panel, right***) demonstration of placement of phantom on driver.

### TBAD Phantom MRE Image Acquisition and Processing

The 4 TBAD phantoms were scanned in a 3T-MR scanner (MAGNETOM Prisma, Siemens Healthcare, Erlangen, Germany). Each phantom was scanned individually and phantoms were positioned in a fixed position with the 10% PVA-c “true lumen” is always facing left and the variable 10%-25% PVA-c “false lumen” facing right. A cardiac-gated spin-echo echo planar imaging (SE-EPI) sequence was used with the following parameters: 5mm thick coronal slices, FOV of 32×32cm^2^, acquisition matrix of 256×256, TR of 228.48ms, TE of 11.93ms, motion encoding gradients (MEG) frequency of 100Hz, and mechanical frequency of 70Hz. Mechanical vibration was introduced into both sides of the phantom using a Resoundant active driver system (Resoundant, Rochester, MN, USA) coupled with an in-house designed passive driver and phantom holder. To better achieve mechanical vibration propagation into the AD phantom, a special pneumatic passive driver and phantom holder were designed and 3D printed.. The phantom housing unit features two rectangular windows 14x36mm (Figure 1 bottom left), to couple pneumatic mechanical vibration generated by the active driver to the phantom with thin diaphragms that vibrate when connected to the active pneumatic driver system. Additionally, the design ensures that the driver produces waves as planar as possible, owing to the 90° angle of vibration relative to the wall of the AD phantom (*12*). The passive driver, as shown in Figure 1, was 3D printed using Stratasys’s VeroClear printing material and a Stratasys Objet 750 Multi Jet 3D printer (Stratasys, Edina, MN, USA).

As this was a validation experiment, phantoms where tested open ended without pressurization of air or fluid within the lumens. Due to the simplicity of the phantom structure and the presence of well-distinguished planar waves, shear stiffness was manually calculated from the wavelength of a planar propagating wave using the equation (Eq.1) *μ* = *ρf*^2^*λ*^2^, where *μ* is the shear stiffness, *ρ* tissue density, *f* external mechanical vibration frequency, and *λ* is the propagating wave wavelength. The wavelength was calculated by counting the number of pixels within one cycle of full wavelength (i.e., red to blue region, Figure 3, annotated by “one cycle”). Then, the number of pixels was converted to spatial displacement by multiplying the number of pixels by pixel size. Finally, the resultant was calculated in Eq.1 as *λ*^2^, 1000g/cm^3^ was used as *ρ*, and 70Hz was used as *f*^2^.

### TBAD Phantom Rheometric Stiffness Testing

Rheometric experiments were conducted on a TA Instruments Discover Hybrid -2 Rheometer (New Castle, DE). 8 mm round samples were cut out of the phantom wall using a punch biopsy blade. The sample is placed on the Peltier plate. For all samples, an 8 mm geometry was lowered to achieve an axial force of 0.300 to 0.340 N with a test spacing of 1mm and a 300 second equilibrium period. Using a previously published method, complex shear modulus was calculated using an oscillation amplitude scan, in deformation control mode, with a strain sweep range of 0.1-20%, at a single frequency of 1 Hz, and a temperature of 25 °C (*13, 14*).

### Healthy Volunteer Thoracic Aorta MRE Image Acquisition and Processing

The thoracic aorta of a 42-year-old male healthy volunteer was scanned with the approval of the institutional review board at the University of Rochester using the same 3T-MR scanner (MAGNETOM Prisma, Siemens Healthcare, Erlangen, Germany). Mechanical vibration was introduced into the thoracic aorta by placing a commercially available passive driver (Resoundant, Rochester, MN, USA) under the volunteer’s left shoulder’s scapular bone in the supine position, as shown in Figure 2. The same SE-EPI sequence was used with the following protocol parameters: 6mm sagittal slices, FOV of 36×36cm2, acquisition matrix of 256×256, TR of 171.36ms, TE of 11.88ms, and eight cardiac segments. Like to the phantom scans, the MEG frequency was 100Hz, and the mechanical frequency was 70Hz. However, MEG encoding was applied in three axes, X, Y, and Z, to encode complex waves that would propagate in all directions.

**Figure 2:**
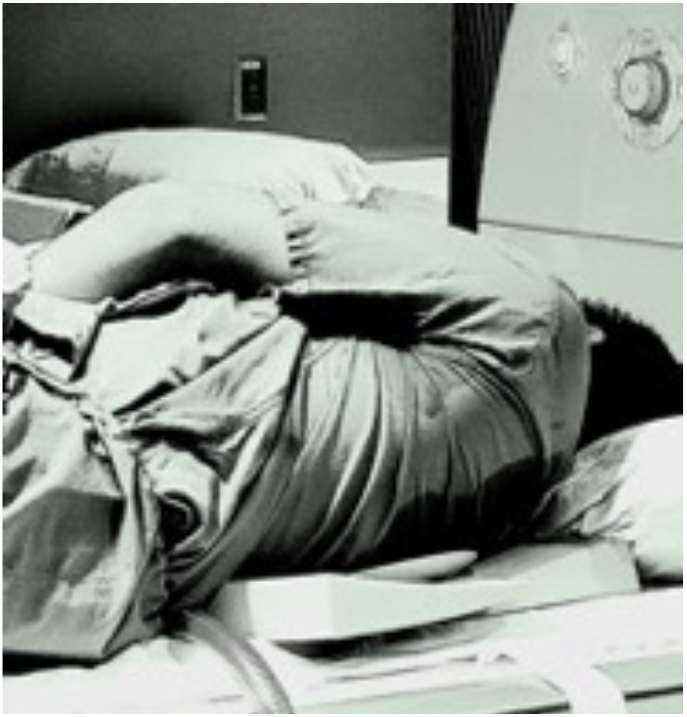
Healthy volunteer setup in which the passive driver (red arrow) is under the scapular bone and the left lateral edge of the vertebral column.

Due to the complexity of the propagating wave within the human body and the difficulty of obtaining clear planar waves, shear stiffness was calculated using the local frequency estimation (LFE) algorithm (MRElab, Mayo Clinic, Rochester, MN) (*12*). First, to minimize blood flow artifacts, the data was acquired in diastole. Second, the thoracic aorta was delineated by an experienced user, marking the region of interest on the magnitude image of the descending thoracic aorta.(Figure 5). Third, an in-plane 8 directions fourth order Butterworth band pass filter was applied to eliminate unwanted longitudinal and reflected waves with cutoff values of 1-40 waves/FOV (*12*). Finally, 3D LFE was used to calculate a single weighted stiffness map. The stiffness map weighting was based on first harmonic amplitude for each encoding direction (i.e. X, Y, and Z)

### Statistical Analysis

We performed a regression model coefficient of determination (R^2^) between rheometric stiffness measurements and MRE generated stiffness estimates, using GraphPad Prism version 10.0.0 for Mac, (GraphPad Software, Boston, Massachusetts).

## Results

### TBAD Phantom MRE and Rheological Stiffness

Figure 3 displays MRE magnitude and wave images of both sides of the TBAD phantoms. Planar waves were clearly observed in the TBAD phantoms (Figure 3). The MRE SE-EPI sequence effectively generated contrasting planar waves in both the false and true lumens of the phantom in the X direction.

**Figure 3:**
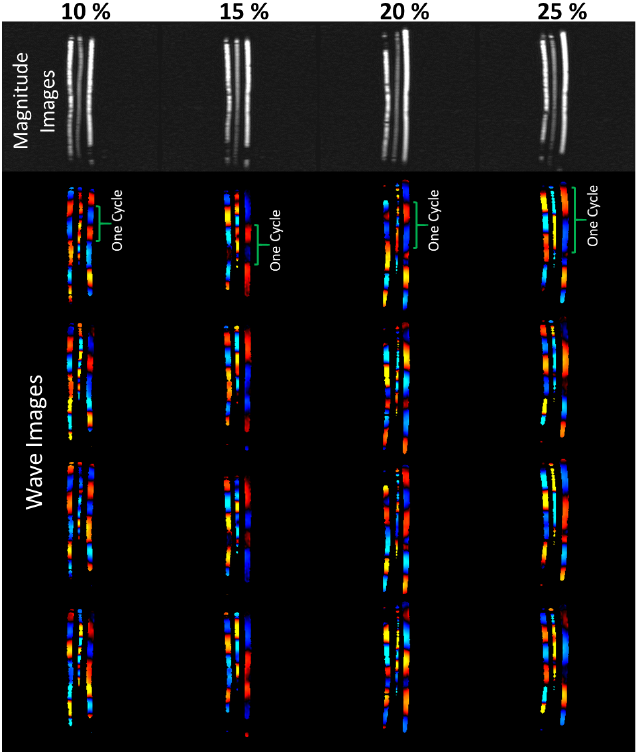
MRE magnitude and wave images of each side of the TBAD phantoms with PVA-c concentrations of 10%, 15%, 20%, and 25%. As shown by the green curled brackets, the wavelength for a given cycle increases with increasing stiffness (i.e. increasing PVA-c concentration).

The wavelength (*μ*) of the 10% PVA-c side of all TBAD phantoms was 81±0.73 mm, corresponding to 64.80±2.68 pixels of size 1.25×1.25 mm, resulting in an MRE shear stiffness of 32.19±2.66 kPa. The wavelength (*μ*) of the 15% PVA-c side of the TBAD phantom was 102.5 mm, corresponding to 82 pixels of size 1.25×1.25 mm, resulting in an MRE shear stiffness of 51.48 kPa. The wavelength (*μ*) of the 20% PVA-c side of the TBAD phantom was 110 mm, corresponding to 88 pixels of size 1.25×1.25 mm, resulting in an MRE shear stiffness of 59.29 kPa. The wavelength (*μ*) of the 25% PVA-c side of the TBAD phantom was 122.5 mm, corresponding to 98 pixels of size 1.25×1.25 mm, resulting in an MRE shear stiffness of 73.53 kPa.

Given the shear rate (frequency) difference between the two modalities (1Hz for Rheologic Testing and 70Hz for MRE) and the viscoelastic properties of the phantoms, the magnitudes of MRE and rheometric-derived stiffness are not expected to be identical. However, the expected increase in stiffness with increasing PVA-c concentration is evident in both modalities (Figure 4). Figure 4 shows the correlation (R^²^ = 0.99) between MRE and rheometric stiffness measurements of the four phantoms with different PVA-c concentrations.

**Figure 4:**
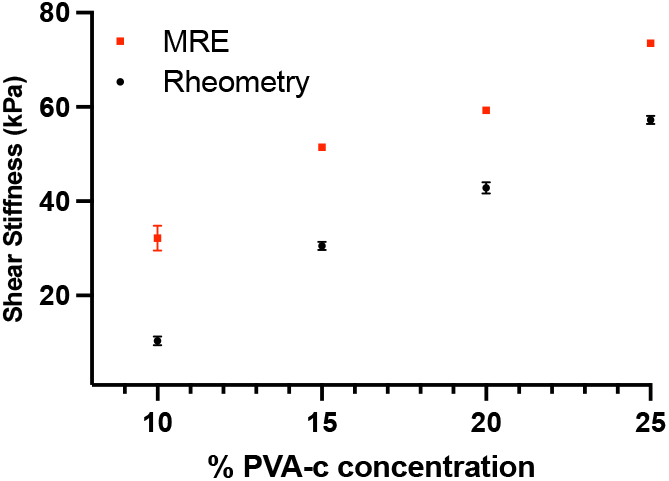
Correlation plot (R^2^=0.99) between MRE and rheometric stiffness measurements of phantoms with different PVA-c concentrations of 10%, 15%, 20, and 25%.

These findings demonstrate the high correlation between rheometric and MRE-generated stiffness results against an independent rheometric gold standard, highlighting the feasibility of using MRE stiffness measurements for TBAD in humans.

### Healthy Volunteer Thoracic Aorta MRE Stiffness

While MRE-derived stiffness measurement to stage liver fibrosis is a clinical routine, the measurement of thoracic aorta mechanical properties has not been previously demonstrated because of known technical difficulties related to motion and flow artifacts by the heart and lungs. Using a novel setup that delivers the mechanical vibration from the anterior side for the subject positioned in supine position, MRE data of a 42 years old male participant was successfully acquired. Figure 5 shows the magnitude and wave images, and stiffness maps of the thoracic aorta of the healthy volunteer. The mean shear stiffness of the thoracic aorta was 3.37±0.83kPa within the aortic ROI and is consistent with previously reported measures of healthy aortic tissue (*8, 15-17*). This demonstrates the capability of performing thoracic aorta shear stiffness measurement with an SE-EPI sequence with the pneumatic MRE driver system with passive driver positioned under the subject’s scapular bone.

**Figure 5:**
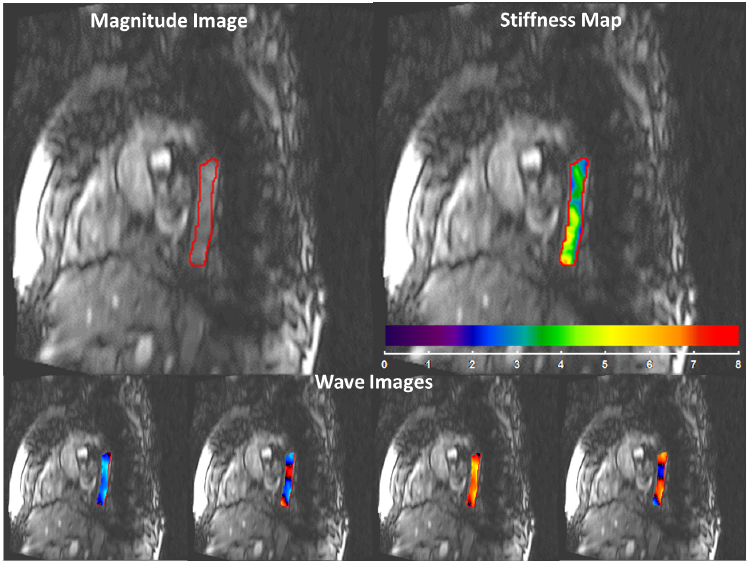
Magnitude image, four snap shots of wave images at different time points in read direction, and stiffness map of the thoracic aorta of a healthy volunteer with a mean shear stiffness of 3.37±0.83kPa.

## Discussion

In this study, we designed and fabricated TBAD phantoms with varying PVA-c concentrations to mimic different stiffness levels in the aortic wall. Our results demonstrate that MRE can effectively capture the stiffness variations across these phantoms, as evidenced by the visualization of planar waves and contrasting stiffness measurements in the false and true lumens (Figure 3). The MRE-derived shear stiffness values for the phantoms with 10%, 15%, 20%, and 25% PVA-c concentrations showed a strong correlation (R^²^ = 0.99) with rheometric measurements, reinforcing the reliability of MRE in capturing biomechanical properties. Furthermore, we successfully scanned and analyzed the thoracic aorta of a healthy volunteer using MRE, demonstrating the feasibility of using MRE to scan the thoracic aorta of a human subject, which is known to be challenging due to blood flow and motion artifacts.

Distinguishing between stable dissections and DAA degeneration remains challenging and affects our understanding and management of TBAD. Although the biomechanical properties of AAA have been extensively profiled (*8, 15-21*), little evidence exists on the evolution of the material properties in DAA over time. In AAA, stiffening of the aortic wall leads to pathological wall stress distribution and predisposes to aneurysmal degeneration and rupture(*4*). Mechanically, increased stiffness due to loss of elastic lamina and increased collagen cross-linking is a significant predictor of aortic dilation (*22-24*). In AD, the medial layer, which is the elastic component of the aorta, is disrupted (*1, 2, 5*) and is expected to make the false lumen consequently stiffer.

The expected difference in stiffness between false and true lumens informs the design of our phantom models. Increased stiffness in the false lumen contributes to elevated pressures (*25*), which are known to drive aneurysmal formation and degeneration through mechanisms such as wall stress(*26*), inflammation, and remodeling (*27*). As such, stiffness indices could become prognostic indicators for DAA formation and growth potentially change clinical decision-making regarding surgical candidates.

The successful application of MRE to measure thoracic aorta stiffness in a healthy volunteer further emphasizes its clinical potential. Prior research on abdominal aortic aneurysms (AAA) has demonstrated the feasibility of quantifying aortic shear stiffness, even in the presence of blood flow and heart pulsation (*16, 17*). Comparisons of aortic MRE shear stiffness estimates with traditional methods, such as pulse wave velocity and 4D flow (*15, 18*), have shown promising results. Additionally, MRE in AAA cases has revealed that stiffness measurements can provide valuable information beyond just aneurysmal diameter (*8*). Consequently, MRE-derived stiffness estimates have the potential to be generalized for use in other aortic regions, including the thoracic aorta, enhancing its utility in various clinical scenarios.

This generalization is supported by studies indicating that MRE can provide valuable insights into vascular health across different aortic segments. Research has shown that MRE-derived stiffness measurements correlate well with traditional metrics like pulse wave velocity (PWV), which is widely recognized for assessing arterial stiffness (*17*). Moreover, MRE offers the advantage of directly visualizing the mechanical properties of the aortic wall, providing a more localized and detailed assessment compared to global measurements like PWV. These capabilities make MRE a promising tool for early detection and monitoring of aortic pathologies, potentially leading to improved patient outcomes through more precise and timely interventions.

## Limitations

Despite the promising results of this study, several limitations should be acknowledged. First, the TBAD phantoms used in this study, while designed to mimic different stiffness levels in the aortic wall, do not fully replicate the complex biological and mechanical environment of a living aorta. The phantoms were fabricated with varying PVA-c concentrations to simulate stiffness, but they lack the dynamic properties and intricate structure of human tissue, potentially limiting the direct applicability of the findings to clinical scenarios. Additionally, while we successfully demonstrated the feasibility of using MRE to scan the human thoracic aorta in a healthy volunteer, blood flow and motion artifacts present significant challenges.

The study’s findings on stiffness differences between false and true lumens in TBAD phantoms are based on controlled experimental conditions. In real-world clinical practice, pathophysiologic variations among patients, including differences in dissection severity, aortic geometry, and individual biological responses, may influence MRE measurements. Therefore, larger studies encompassing diverse patient profiles are required to translate these findings comprehensively.

Finally, sample size is a limitation of our study. The successful application of MRE in thoracic aorta was demonstrated in only one healthy volunteer. Studies with more extensive and diverse subject populations are essential to establish the clinical utility of MRE in assessing thoracic aorta stiffness. Including a broader range of subjects, such as those with different stages of aortic diseases and varying demographic characteristics, will provide more robust data and help validate the findings across a broader spectrum of clinical scenarios.

## Conclusion

This study demonstrates excellent agreement between stiffness measurements obtained using MRE and rheometry in TBAD phantoms. In addition, a feasibility study was performed in a healthy volunteer to determine the stiffness of thoracic aorta despite of heart and lung motion. We also present a method for the novel manufacturing of hydrogel aortic dissection phantoms with programmatic regional stiffness variations measured using MRE and validated against Rheometric measurements.

